# GILEA: GAN Inversion-enabled latent eigenvalue analysis for phenome profiling and editing

**DOI:** 10.1101/2023.02.10.528026

**Authors:** Jiqing Wu, Viktor H. Koelzer

## Abstract

Modeling heterogeneous disease states by data-driven methods has great potential to advance biomedical research. However, a comprehensive analysis of phenotypic heterogeneity is often challenged by the complex nature of biomedical datasets and emerging imaging methodologies. Here, we propose a novel GAN Inversion-enabled Latent Eigenvalue Analysis (GILEA) framework and apply it to phenome profiling and editing. As key use cases for fluorescence and natural imaging, we demonstrate the power of GILEA using publicly available SARS-CoV-2 datasets stained with the multiplexed fluorescence cell-painting protocol as well as real-world medical images of common skin lesions captured by dermoscopy. The quantitative results of GILEA can be biologically supported by editing latent representations and simulating dynamic phenotype transitions between physiological and pathological states. In conclusion, GILEA represents a new and broadly applicable approach to the quantitative and interpretable analysis of biomedical image data. The GILEA code and video demos are publicly available at https://github.com/CTPLab/GILEA.

## Introduction

Phenomics^1^ - the systematic study of traits that make up a phenotype - is a main driver of scientific insights into human physiology and the pathogenesis of diseases. Recent progress in this field has been driven by the emergence of new imaging technologies, enabling the high-content analysis of disease conditions ^2,3^. However, an in-depth analysis of phenotypic heterogeneity and dynamic transitions between physiological and pathological states is often challenged by the static and complex nature of biomedical imaging datasets. There is a clear need for a robust and comprehensive feature quantification methodology applicable to large biomedical datasets derived from both established and emerging imaging technologies. Further, it is of key interest to generate in-silico predictions of dynamic transitions between biological states. As most biomedical imaging methods capture only a snapshot of a given biological system, current approaches for phenomic profiling ^4,5^ are constrained by static-imaging-based models and fail to explore dynamic and complex phenotypic changes that underlie phenotype heterogeneity. Recently, the study of GAN Inversion^6,7^ has brought novel insights into understanding and controlling latent semantic representations. Thanks to the learned representations of real images that lie in the latent space of GAN, researchers can search for latent vectors of interest and edit the attributes of real images. Considering biomedical imaging as a specific instance of the image domain, we are motivated to apply **G**AN **I**nversion-enabled **L**atent **E**igenvalue **A**nalysis (GILEA) for profiling and editing latent representations of phenotypic image data.

### GILEA

In this article, we apply GILEA to two widely-used biomedical imaging modalities: fluorescence and natural imaging. First, we investigate GILEA on two large-scale fluorescence microscopy SARS-CoV-2 datasets RxRx19 (a, b)^5^ for cell-based drug screening. Following the multiplexed cell-painting protocol^8^, these datasets allow to reveal relevant cellular components and organelles (e.g., Nucleus (DNA), Endoplasmic reticulum (ER), Actin (Actin), Nucleolus and cytoplasmic RNA (RNA), Golgi and plasma membrane (Golgi)) in a highly multiplexed, high-content manner. With the inclusion of more than 1800 drug compounds with up to 8 different concentrations that are tested on 3 different cell-lines, the main experiments of GILEA are carried out in a phenotypic drug effects analysis. Second, we demonstrate the domain-agnostic application of GILEA to the natural image domain, represented by the multi-source HAM10000 dataset of more than 10000 dermoscopic images of common pigmented skin lesions ^9^. In both settings, we conduct thorough ablative studies, compare *d*_LEA_ to existing statistical measurements and verify the performance under different GAN inversion architectures.

To measure the difference between two collections of (natural) images, researchers have proposed a variety of evaluation methods such as Fréchet Inception Distance (*d*_FID_)^10^, Inception Score^11^ and Kernel Inception Distance (*d*_KID_)^12^. These approaches could be used to differentiate image feature heterogeneity between disease states in a given biomedical dataset, but it is non-trivial to derive an in-depth understanding of biological states with these scores and to support research insights with plausible visual interpretations. Therefore, prior art is less satisfactory for critical biomedical applications. Motivated by the emergence of (unsorted) eigenvalues in the improved implementation of *d*_FID_, we recently ^13^ suggested comparing sorted eigenvalues (*d*_Eig_) as a simple alternative to *d*_FID_. For *i* = 1,2, consider 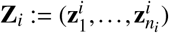 be a collection of *n_i_ p*-dimensional vectors, we have

#### Definition 1.

*Let* 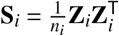 *be the sample covariance matrix (SCM) of* **Z**_*i*_., *we define*

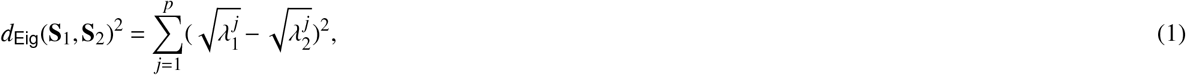

*where* 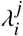 *is the j-th largest eigenvalue of* **S**_*i*_.

#### Quantification of phenotypic heterogeneity

In the study of *d*_Eig_, **Z**_*i*_ is usually the collection of features obtained with the penultimate layer (*pool3*) of an Inception V3 model^14^, where the model is trained for an ImageNet classification task. However, such an Inception model trained with ImageNet is not suitable for deriving biological-aware representations. Alternatively, we utilize GAN Inversion^6^ and propose to learn the latent representations **Z**_*i,k*_ of individual cellular organelles (Fig. 1 (b)) or tissue components (‘Methods’ section). To address the infeasible deployment of *d*_Eig_ to cases where we obtain imbalanced values of *d*_Eig_ for *k* = 1,…, *c*, we propose

#### Definition 2.

*For i* = 1,2 *and k* = 1,…,*c, let* 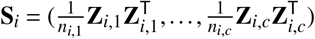 *be the collection of SCMs of **Z**_i,k_, then we define*

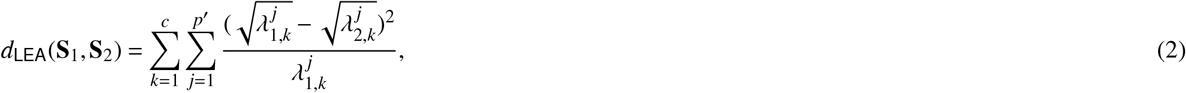

*where p*′ ≪ *p and* **S**_1_ *is the reference SCM.*

**Figure 1.**
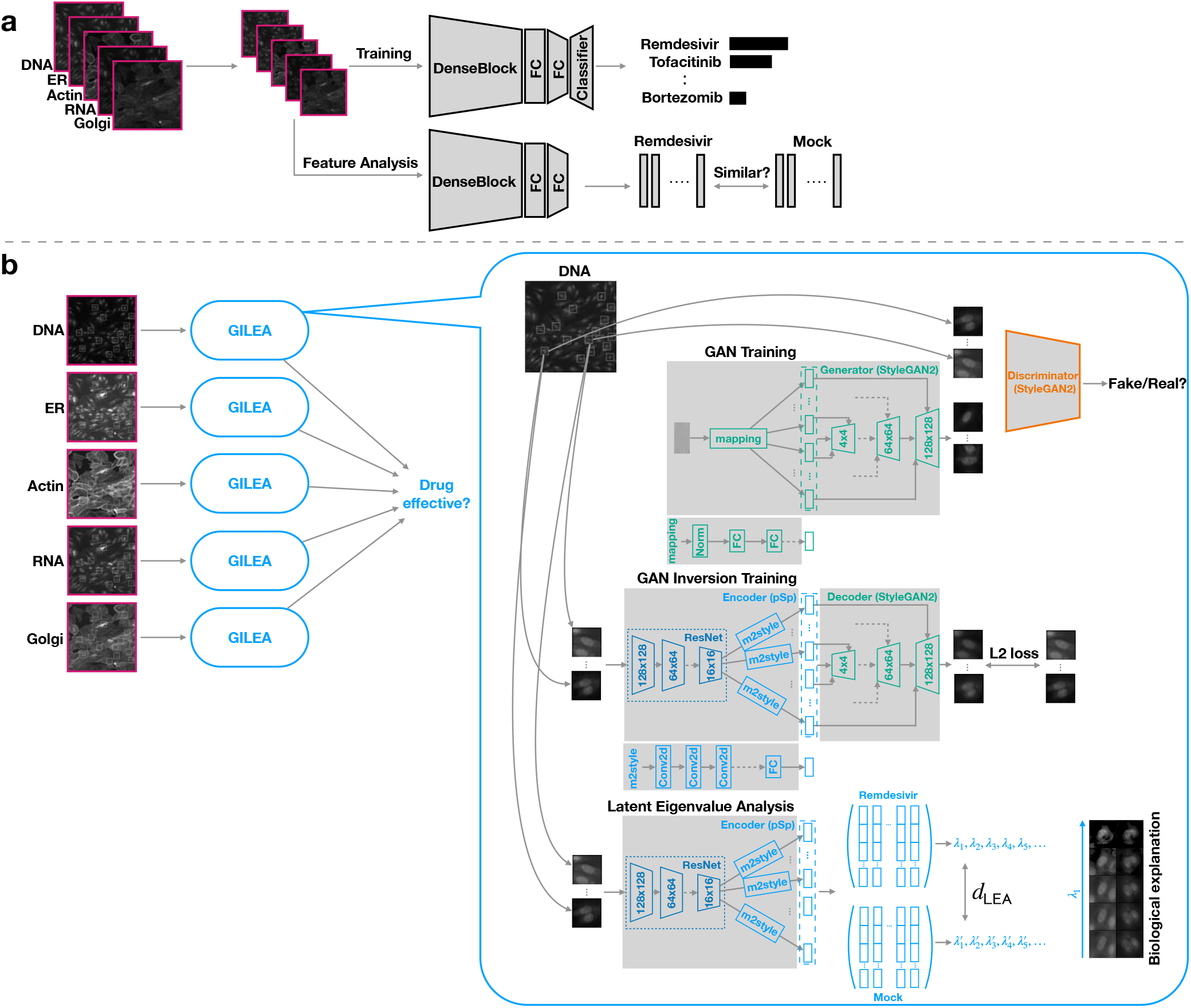
Model illustrations for the baseline method (Cuccarese *et al.* ^5^) **(a) and proposed GILEA approach (b)**. **a**, In the baseline method, a variant model of DenseNet-161 is trained on the downsized multiplexed fluorescent images for classifying the drug compounds (Remdesivir, Tofacitinib, Bortezomib shown as exemplars). Then, the learned features are utilized to analyze the effectiveness of different drugs. **b**, In GILEA, we start the pipeline by pre-training the decoder on center-cropped single-cell images in an adversarial training manner for each fluorescent channel. Then, we learn robust latent representations with a residual-based encoder^20^ for reconstructing these single-cell images. For quantifying the drug effects, we compute the eigenvalues with learned single-cell representations and support our quantitative results with clearly observable phenotypic transitions.

Similar to Principal Component Analysis (PCA)^15^, we only utilize the *p*′ ≪ *p* largest eigenvalues that reflect the largest variances and the most critical information. As the 5 largest eigenvalues dominate > 95% of the overall values in the experiments, we set *p*′ = 5 throughout the article (please see ‘Results’ and ‘Methods’ for more detail).

#### Visualization of phenotypic transitions

Complementary to the *d*_LEA_ that measures the eigenvalue difference along each principal axis, we simulate observable phenotypic transitions by editing latent representations. Concretely, this is achieved via controlling their principal component(s), which enables a direct linkage of quantifying phenotypic heterogeneity and presenting plausible phenotypic changes. Notably, this linkage between quantification with the largest eigenvalues and visualization with their principle components is missing for most existing studies in latent semantic understanding of (fake) natural images^16–19^. For example, Härkönen *et al.* ^17^ proposed to edit fake images by adding weighted eigenvectors to its latent representations. Similarly, Shen *et al.* ^16^ utilized the closed-form factorization to determine the manipulation direction. Since such manipulation directions are not parallel to the principal axes, they cannot be directly used to explain the eigenvalue heterogeneity embedded in *d*_LEA_. To resolve this gap, we propose to output an image sequence by manipulating the largest principal component(s) of latent representations **Z**_*i,k*_ (Def. 2), where **Z**_*i,k*_ are derived from real biomedical images.

#### Definition 3.

*Following the specifications of* 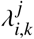, **Z**_*i,k*_ *of Def. 2, let* 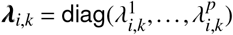 *be decomposed as* 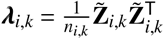, *where* 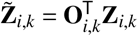 *are the principal components w.r.t. the orthogonal eigenbasis* **O**_*i,k*_. *Given j* ∈ {1,…, *p*′}, *we interpret the difference between* 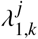 *and* 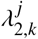 *(Eq. 2) by visualizing* 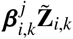 *manipulated on the *j*-th principal component that brings*

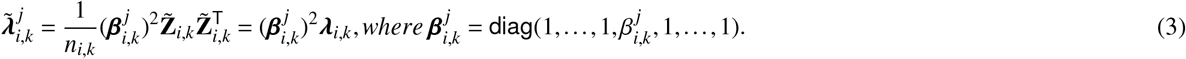

To guarantee the consistency and to simulate clearly interpretable phenotypic transitions, image sequences *w.r.t*. 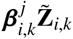 are always obtained by step-wise assigning 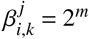 in this study, where *m* = (−1.8, −1.2, −0.6,0,0.6,1.2,1.8). In summary, our contributions can be summarized as follows:

- Built on the promising GAN Inversion framework, we learn rich latent representations of biomedical image data that enable the fine-grained analysis for phenome profiling and editing.
- By comparing the largest eigenvalues *d*_Eig_, we propose the numerically robust quantification *d*_LEA_ of phenotypic heterogeneity. By controlling the largest principal components, we edit the latent representations and simulate phenotypic transitions that deliver plausible interpretations of *d*_LEA_.
- We demonstrate the strength of GILEA by application on two large-scale SARS-CoV-2 drug screening datasets imaged with the high-content cell painting protocol. Further, we illustrate the domain-agnostic facet of GILEA by application to natural images derived from clinical workflows and confirm its wide applicability in biomedical research.

## Results

Here, we report the application of GILEA to two large-scale fluorescence microscopy datasets RxRx19 (a,b) released by Recursion^5^, which document the phenotypic effects of more than 1800 drug candidates on Severe Acute Respiratory Syndrome Coronavirus Type 2 (SARS-CoV-2) infection and associated systemic inflammation using the multiplexed cell-painting protocol on human and animal cell-lines. We set the drug hit score proposed by Cuccarese *et al.* ^5^ as the baseline and demonstrate the performance of *d*_LEA_. To investigate the phenotypic effects of drug candidates at single-cell resolution, we carried out cell segmentation using the DNA channel (Please see the Mahotas documentation^21^ or our code repository). Accordingly, we derive 23 million 64 × 64 single-cell training images from 0.37 million raw images. This allows us to analyze drug effects on the individual cell organelle, greatly extending the range of detectable phenotypic perturbations. As shown in Fig. 1 (a,b), our approach using unsupervised training on the center-cropped cell images for the individual fluorescent channels indeed differs greatly from the published baseline using supervised training on the entire image read-outs. As the phenotypic library (RxRx19a) has four cell conditions: Mock control (Mock), irradiated control (Irradiated), infected without drug treatment (Infected), and infected with different drug treatments (Drug), we set the reference **S**_1=Mock_ (Eq. 2) corresponding to the ‘Mock’ latent representations and then determine the effect of a drug based on whether it reverses the phenoprint of infected cells. *d*_LEA_ is provided as percentage (×100) in the following plots for clearer visualization.

#### Definition 4.

*Following the specification ofSCMs* **S**_Mock_, **S**_Infected_, **S**_Drug_ *as Eq. 2, we define*

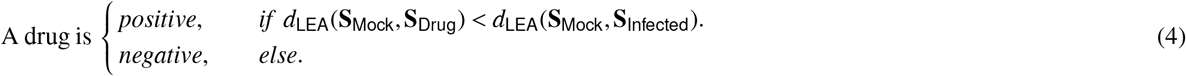

**Table 1.** The numerical comparison of reconstruction results between the proposed and ensemble GILEA on VERO and HRCE.

|  |  | VERO (Proposed) | VERO (Ensemble) | HRCE (Proposed) | HRCE (Ensemble) |
| --- | --- | --- | --- | --- | --- |
| DNA | PSNR | $35.24 \pm 1.87$ | $34.67 \pm 2.29$ | $44.32 \pm 2.48$ | $38.25 \pm 3.66$ |
| | SSIM | $0.95 \pm 0.01$ | $0.95 \pm 0.02$ | $0.99 \pm 0.01$ | $0.95 \pm 0.05$ |
| ER | PSNR | $35.31 \pm 1.94$ | $32.20 \pm 2.11$ | $35.27 \pm 2.36$ | $29.42 \pm 3.12$ |
| | SSIM | $0.96 \pm 0.01$ | $0.93 \pm 0.02$ | $0.95 \pm 0.02$ | $0.83 \pm 0.09$ |
| Actin | PSNR | $33.54 \pm 2.70$ | $31.74 \pm 2.72$ | $33.68 \pm 3.21$ | $29.22 \pm 3.44$ |
| | SSIM | $0.93 \pm 0.02$ | $0.90 \pm 0.03$ | $0.92 \pm 0.04$ | $0.78 \pm 0.09$ |
| RNA | PSNR | $36.93 \pm 1.51$ | $34.59 \pm 1.87$ | $39.00 \pm 2.56$ | $33.87 \pm 3.07$ |
| | SSIM | $0.97 \pm 0.01$ | $0.95 \pm 0.02$ | $0.96 \pm 0.02$ | $0.90 \pm 0.06$ |
| Golgi | PSNR | $37.23 \pm 2.24$ | $33.84 \pm 2.08$ | $36.48 \pm 2.44$ | $32.69 \pm 3.29$ |
| | SSIM | $0.95 \pm 0.02$ | $0.92 \pm 0.02$ | $0.95 \pm 0.03$ | $0.87 \pm 0.07$ |
| Total | PSNR | $35.27 \pm 1.86$ | $33.14 \pm 2.11$ | $36.26 \pm 2.40$ | $31.36 \pm 2.95$ |
| | SSIM | $0.95 \pm 0.01$ | $0.93 \pm 0.02$ | $0.95 \pm 0.02$ | $0.87 \pm 0.07$ |

### GILEA benchmarking

For experiments carried out on both human (Human renal cortical epithelial cells, HRCE) and animal (Kidney epithelial cells of African green monkey, VERO) cell-lines, we start our investigation by comparing the reconstruction quality between the proposed GILEA – trained on individual fluorescent channels and ensemble GILEA – trained on all the channels. For each fluorescent channel, the proposed GILEA outperforms the ensemble variant on both cell-line experiments in terms of overall better Peak signal-to-noise Ratio (PSNR) and Structural Similarity Index Measure (SSIM) scores (Tab. 1). This indicates superior reconstruction quality by the proposed GILEA over the ensemble approach (Please see Fig. 2 (a) and (f) for visual results). Importantly, the proposed GILEA is well calibrated by the small distance between mock control and irradiated control cells (Fig. 2 (b,d,g,i)), which is consistent with the expected similar phenotypic and biological characteristics shared by the control conditions ‘mock’ (cells in culture medium without viral stimulation) and ‘irradiated’ (cells in culture medium incubated with the inactivated virus). In contrast, such a verifiable calibration cannot be reproduced with the ensemble approach. For the HRCE experiment, Fig. 6 (c, d) in the Appendix shows an inexplicably small difference between mock and infected cell populations as compared to the control conditions, which contradicts the expected phenoprint heterogeneity induced by the virus. In terms of the largest eigenvalues, *p*′ = 5 robustifies the drug effect quantification of *d*_LEA_ and shows an improved consistency with the hit score^5^, which clearly differs from *p*′ = 1, a balanced alternative of *d*_Eig_ (See Fig. 7 (VERO) and 8 (HRCE) in the Appendix for more detail).

**Figure 2.**
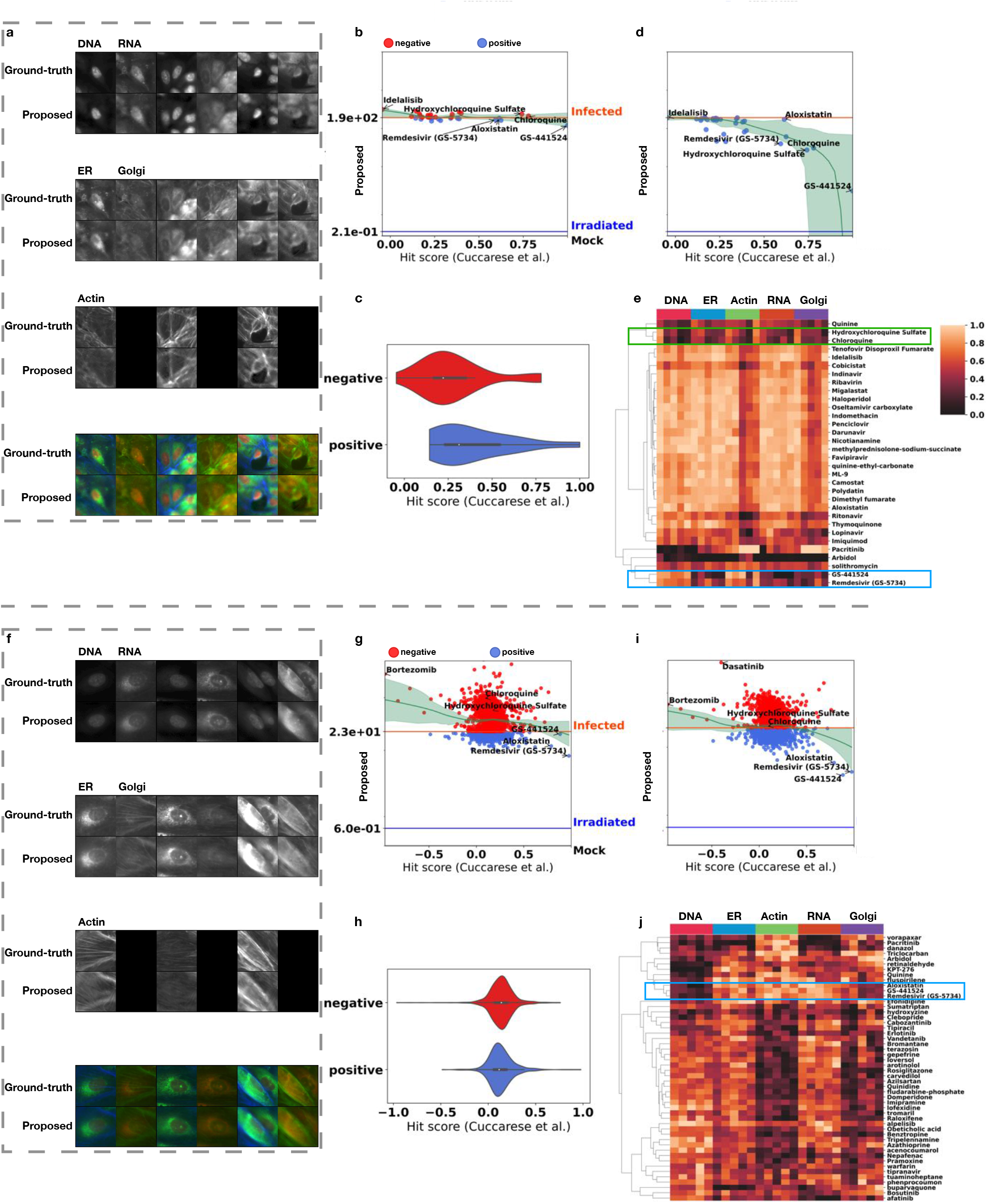
Reconstruction visualization of GILEA and quantitative comparison of drug responses between the baseline (Cuccarese *et al.* ^5^) and *d*_LEA_ (Proposed). **a** (VERO) and **f** (HRCE): The reconstructed samples obtained by GILEA. **b** (VERO) and **g** (HRCE): The quantitative comparison between the hit score^5^ and *d*_LEA_ with the latent representations of all concentrations. **c** (VERO) and **h** (HRCE): The violin plot of overall comparison between the hit score^5^ and *d*_LEA_. **d** (VERO) and **i** (HRCE): The quantitative comparison between the hit score^5^ and dLEA with the latent representations of optimal drug concentration. **e** (VERO) and **j** (HRCE): The hierarchical clustering of top 50 drug compounds (if exist) *w.r.t.* the 5 largest eigenvalues of the latent representations of optimal drug concentration.

### Overall comparison

For a thorough comparison, we screened all drug compounds tested on VERO and HRCE cell-lines (with the exclusion of three drugs that have duplicated or ambiguous names). Despite fundamentally different model designs (Fig. 1 (a,b)), our quantitative score *d*_LEA_ demonstrates an overall consistent correlation to the baseline hit score^5^, that is, the lower the *d*_LEA_ is, the higher the baseline hit score is. Importantly, the effect estimation for a given drug compound can be directly derived from *d*_LEA_, while a manual threshold determination is required for the hit score (baseline) ^5^. As displayed in Fig. 2 (b, c) of the VERO experiments, *d*_LEA_ shows a mild yet meaningful decreasing trend with growing hit scores and identifies Remdesivir and its prodrug GS-441524 as efficacious compounds when using all their latent representations that are independent of concentration. Further, GILEA allows to take the optimal drug concentration into consideration, and thus achieves a superior resolution in identifying effective drug candidates as illustrated in Fig. 2 (d). Importantly, Remdesivir and GS-441524 remain the top effective candidates among all the drug compounds. The superior effectiveness of Remdesivir and GS-441524 can also be differentiated from other drugs by examining the hierarchical clustering (Fig. 2 (e)). For example, we observe distinct patterns of latent representations of ER, RNA and Golgi channels for Remdesivir and GS-441524. This corresponds to the successfully reversed phenoprint reflected by the largest eigenvalues, *e.g.,* (ER ×10^5^): 1.33 for Mock, 1.33 for GS-441524, 1.36 for Remdesivir, and 1.66 for Infected cells. (RNA ×10^5^): 1.75 for Mock, 1.76 for GS-441524, 1.79 for Remdesivir, and 2.20 for Infected cells. (Golgi ×10^5^): 3.23 for Mock, 3.20 for GS-441524, 3.23 for Remdesivir, and 3.69 for Infected cells.

Similar to the VERO experiment, Fig. 2 (g) and (h) show that *d*_LEA_ remain well correlated with the baseline in the HRCE experiment. Strikingly, Remdesivir and GS-441524 are identified as strongly efficacious when computing the eigenvalues on the latent representations of the optimal drug concentration (Fig. 2 (i)), indicating verifiable positive drug effects achieved by both candidates. On the other hand, Chloroquine and Hydroxychloroquine demonstrate contradictory effects on both the latent representations of the cells treated with different drug concentrations and the optimal drug concentration, both of which are identified as negative by the *d*_LEA_(**S**_Mock_, **S**_Infected_) threshold (Fig. 2 (g) and (i)). Such inconsistency between the ineffective identification on the human cell-line and the effective identification on the animal cell-line undermines its fidelity in clinical treatment, which can be explained by the fact that neither of them is recommended in treating hospitalized COVID-19 patients according to clinical studies^22,23^. If we examine the latent representations in more detail, Fig. 2 (j) highlights the unique patterns presented in ER, Actin, and RNA channels for Remdesivir and GS-441524, which reveal novel and subtle phenotypic changes that were previously unidentified. Cell-level phenotype analysis by *d*LEA can therefore provide novel insights and patterns in high-throughput drug-screening experiments as candidates for subsequent biological exploration.

### Fine-grained quantification and visual interpretation

Now, we report the fine-grained quantification on both VERO and HRCE experiments for individual fluorescent channels and drug concentration levels. As for the overall analysis, the small difference between mock and irradiated control is correctly captured in individual fluorescent channels for both experiments, which serves as an important sanity check for the stratified quantification. Regarding the drugs of interest presented in Fig. 3 (a) and Fig. 4 (a), we observe a consistently improved phenoprint rescue with an increasing dose level of the drug. This reflects the biologically plausible observation of increased inhibitory effects on cellular infection by SARS-CoV-2 with increasing drug concentrations. For VERO and HRCE cell-lines, Remdesivir and GS-441524 are consistently identified as effective compounds *w.r.t.* individual fluorescent channels as well as all channels, which supports the clinical utility of Remdesivir approved by U.S. Food and Drug Administration (FDA)^1^. Meanwhile, the heterogeneous effects of Chloroquine and Hydroxychloroquine are also revealed in our refined analysis. For instance, the negative hit results in Actin, RNA and Golgi channels eventually undermine the overall performance of both candidates and provide the negative evidence for both drugs derived from real-world clinical studies^24^. Furthermore, the PCA plots displayed in (b) of Fig. 3 and 4 clearly support the efficacious hit results achieved by Remdesivir. With the k-means clustering on the largest principal component(s), we observe meaningful groups based on the nucleus morphology (DNA) for both VERO and HRCE. Besides, interesting and striking phenotypic transitions arise in other understudied channels. Taking the RNA channel as a concrete example, the largest eigenvalue (×10^5^) of mock and infected cells are 1.75, 1.23 versus 2.20, 1.28 for VERO and HRCE *resp.* As shown in (b) of Fig. 3 and 4, the cellular sequences presented from left to right with enlarging the largest principal component(s) imply increased RNA production in the cytoplasm. This observation is biologically plausible, as SARS-CoV-2 expresses RNA-dependent RNA polymerase as well as a large number of supporting factors to transcribe and replicate the viral genome in infected cells^25^. Viral infection of host cells thus leads to massive upregulation of the production of viral RNA in the cytoplasm, which is correctly identified by the *d*_LEA_ analysis at subcellular resolution. Importantly, Remdesivir acts as a nucleoside analog and stalls the RNA-dependent RNA polymerase of coronaviruses^26^. Our analysis identifies this effect at the phenotypic level, as Remdesivir treatment calms the hyper state reflected by shifting the largest eigenvalue from 2.20 to 1.79 (VERO) and from 1.28 to 1.26 (HRCE). Taking the 5 largest eigenvalues as a whole, we further robustify and differentiate the positive drugs from negative ones, while demonstrating persistent transitions for all fluorescent channels.

**Figure 3.**
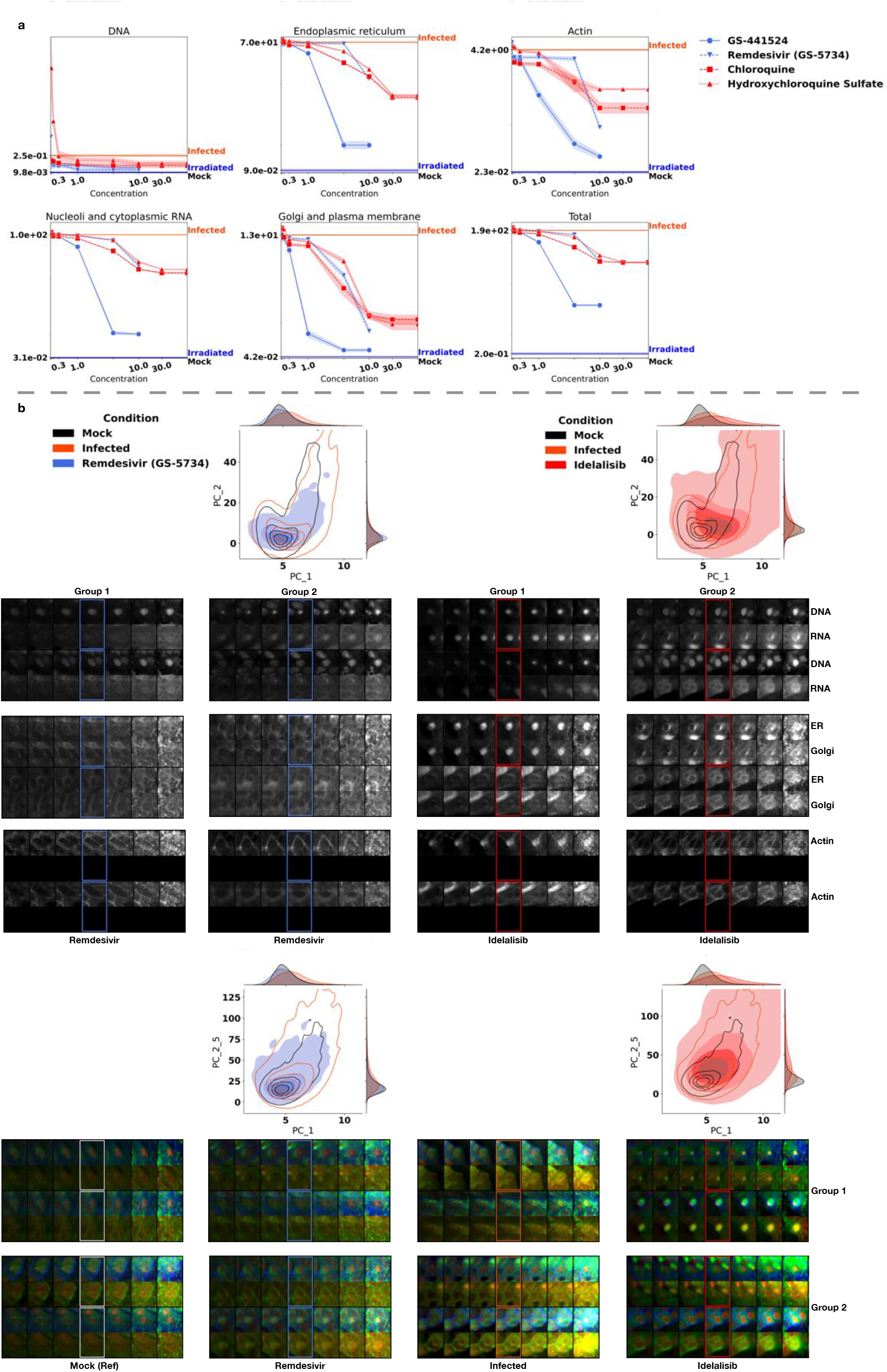
Identification of drug-concentration dependent effects and visual interpretation for key drugs of interest in the VERO cell-line. **a**, The proposed *d*_LEA_ of different drug concentrations for individual and all fluorescent channels. Here, we report the mean *d*_LEA_ (with standard deviation) averaged on 4 randomly sampled cell collections. **b**, The PCA plots and phenotypic transitions driven by manipulating the largest (top) and 5 largest (bottom) principal component(s). The bounding box indicates the reconstructed image.

**Figure 4.**
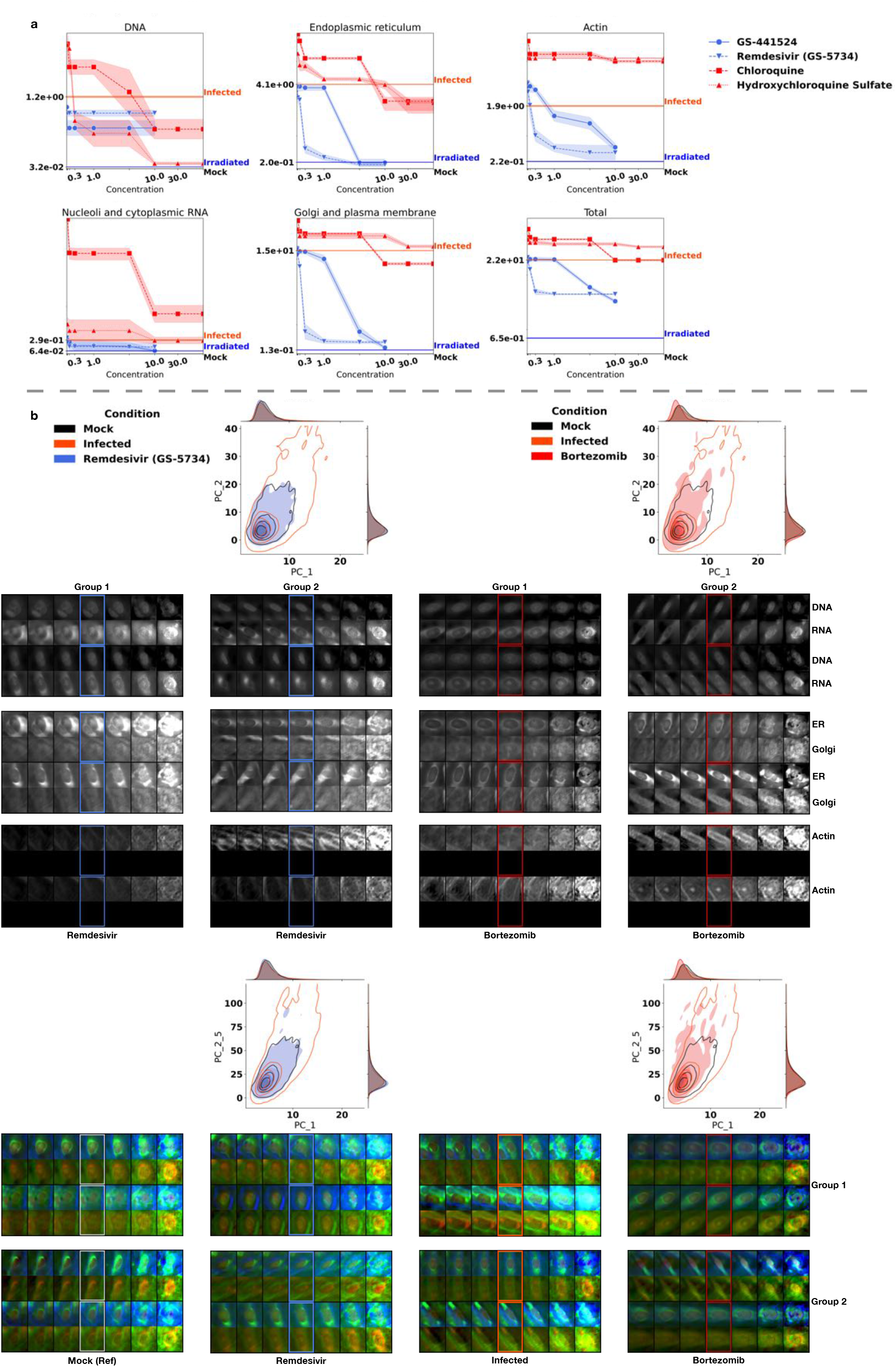
Identification of drug-concentration dependent effects and visual interpretation for key drugs of interest in the HRCE cell-line. **a**, The proposed *d*_LEA_ of different drug concentrations for individual and all fluorescent channels. Here, we report the mean *d*_LEA_ (with standard deviation) averaged on 4 randomly sampled cell collections. **b**, The PCA plots and phenotypic transitions driven by manipulating the largest (top) and 5 largest (bottom) principal component(s). The bounding box indicates the reconstructed image.

## Discussion

In this study, we propose the novel GILEA approach for profiling and editing latent phenotypic representations of biomedical imaging data. We demonstrate the practical application of GILEA on the multiplexed fluorescence microscopy SARS-CoV-2 datasets at single-cell and single-organelle granularity. Through application to well-defined biological settings, we achieve refinement of high-throughput drug screening under the cell-painting protocol. Importantly, we verify the GILEA results in direct comparison to the baseline method^5^, which is implemented with a fundamentally different approach. The drug candidates of interest learned from GILEA in the animal (VERO) and human (HRCE) SARS-CoV-2 infection experiments can be plausibly supported by prior medical knowledge and thus suggest the plausible and effective identification of drug effects using GILEA. At the same time, we acknowledge the limitation of GILEA on the human umbilical vein endothelial (HUVEC) cell-line, which models cytokine storm conditions in severe COVID-19^27^. Although the least effective drug candidates identified by Cuccarese *et al.* ^5^ are likely to be assigned with high *d*_LEA_ score (Appendix Fig. 9), the association is less clear for drug candidates with a positive baseline hit score (*e.g.,* c-MET inhibitors in Fig. 9 (b)). This suggests that the current cell-level evidence is not sufficient and follow-up in-vivo studies are needed.

Importantly, GILEA is not limited to multiplexed fluorescence imaging as the input modality. In a different biomedical use case, we conduct thorough ablative studies on GAN Inversion model architectures and apply GILEA to the natural imaging dataset HAM10000^9^. This dataset represents multi-source dermatoscopic images of common pigmented skin lesions, and is representative of a common ‘clinical grade’ imaging modality. In application to HAM10000 we demonstrate the quantification of phenotypic heterogeneity across different lesion categories and investigate human-interpretable phenotypic transitions. From a domain perspective, our approach thus provides new possibilities for the high-throughput analysis of biomedical image datasets, and for the re-interpretation of feature heterogeneity in a biomedical context. These use cases highlight the utility of GILEA to enable virtual ‘phenome editing’ at the interface of advances in GAN Inversion and biomedical imaging approaches.

Conceptually, this approach has the potential to open a completely new path toward AI-enabled simulation of biological dynamics in a wide range of physiological and pathological settings. Specifically, learned representations of biomedical systems could be used to simulate diverse pathological and physiological conditions as well as matched therapeutic interventions. Following the algorithmic ‘phenome editing’, an important future study is the extension of GILEA to in-silico ‘genome editing’ that drives and controls cellular changes. With reference to the labor-intensive and time-consuming biological editing, a GAN Inversion-enabled computational ‘scissor’ will allow high-throughput and selective computational interventions on (latent) genetic representations and may therefore serve as a promising proxy tool for targeted disease therapeutics.

## Methods

In reference to Fig. 1(b), here we describe the architecture of GILEA in detail and provide insights into the evaluation of model performance. This is done by conducting a separate series of ablative experiments on the natural imaging dataset HAM10000, a large collection of multi-source dermatoscopic images of common pigmented skin lesions^9^. Complementary to above drug screening studies, these experiments support the technical robustness of the GILEA pipeline to various types of input data and showcase its domain-agnostic facet. As HAM10000 includes clinical images of human skin lesions, it is easily interpretable by domain experts and allows conceptual validation of the GILEA results in the context of well-established disease categories. Concretely, the HAM10000^9^ dataset has 7 categories of skin lesion images including actinic keratoses (akiec), basal cell carcinoma (bcc), benign keratosis (bkl), dermatofibroma (df), melanocytic nevi (nv), melanoma (mel) and vascular skin lesion (vasc). As the RGB channels of skin lesion images jointly inform the clinical presentation, we train the GILEA pipeline on images with all RGB channels simultaneously. For the sake of probing *d*_LEA_ within such a distinct domain, we design simple interpolation experiments among different categories as follows. Considering the benign nevus (‘nv’) collection of images as the reference and comparing this to the malignant counterpart (e.g., malignant melanoma, mel), we randomly mix the images of ‘nv’ and ‘mel’ (e.g., **x**_mel,1_,…,**x**_mel,*n*_mel__,**x**_nv,1_,…,**x**_nv,*n*_v__) according to the interpolation weight 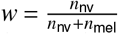. Then, we measure the distribution difference between nv and the mixed collection by *d*_LEA_. Ideally, we should observe that *d*_LEA_ converges to 0 when w is shifted from 0 to 1 with the increasing inclusion of ‘nv’ (*n*_nv_ ↑) and exclusion of ‘mel’ images (*n*_mel_ ↓) (Fig. 5 (b)). To avoid the sample imbalance between the reference (nv, 6705 images) and compared categories, we take mel (1113), bkl (1099), and bcc (512) for comparison.

**Figure 5.**
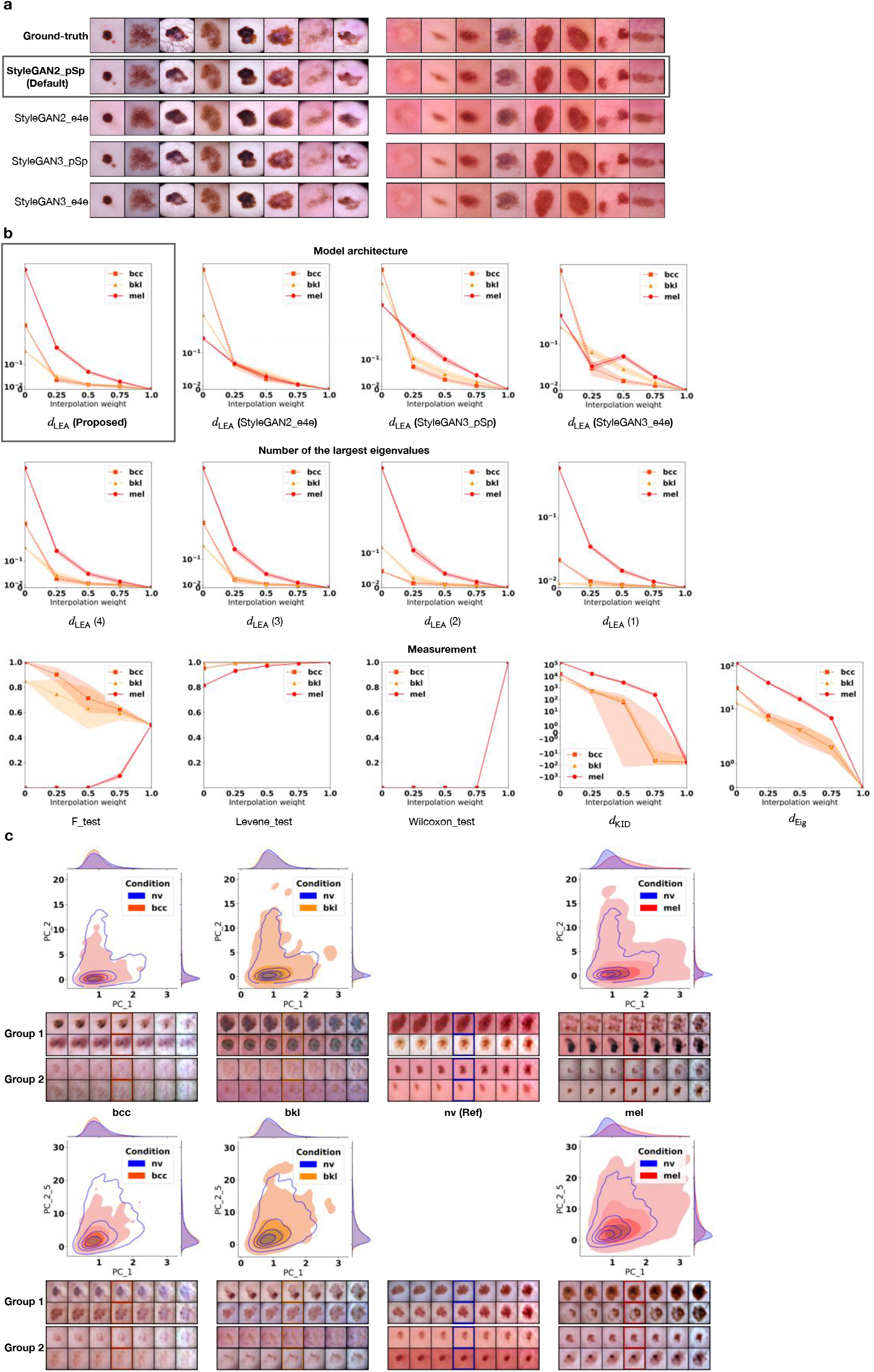
The data interpolation quantification and visual interpretation on HAM10000 experiments. **a**, The reconstructed samples obtained by 4 different architectures. **b**, The *d*_LEA_ comparison of data interpolation regarding different architectures, number of the largest eigenvalues, and existing measurements. Here, we report the mean *d*_LEA_ and comparative measurements (with standard deviation) averaged on 4 randomly sampled data mixtures given the interpolation weight. **c**, The PCA plots and morphological transitions driven by manipulating the largest principal components. The bounding box respectively indicates the reconstructed image.

**Figure 6.**
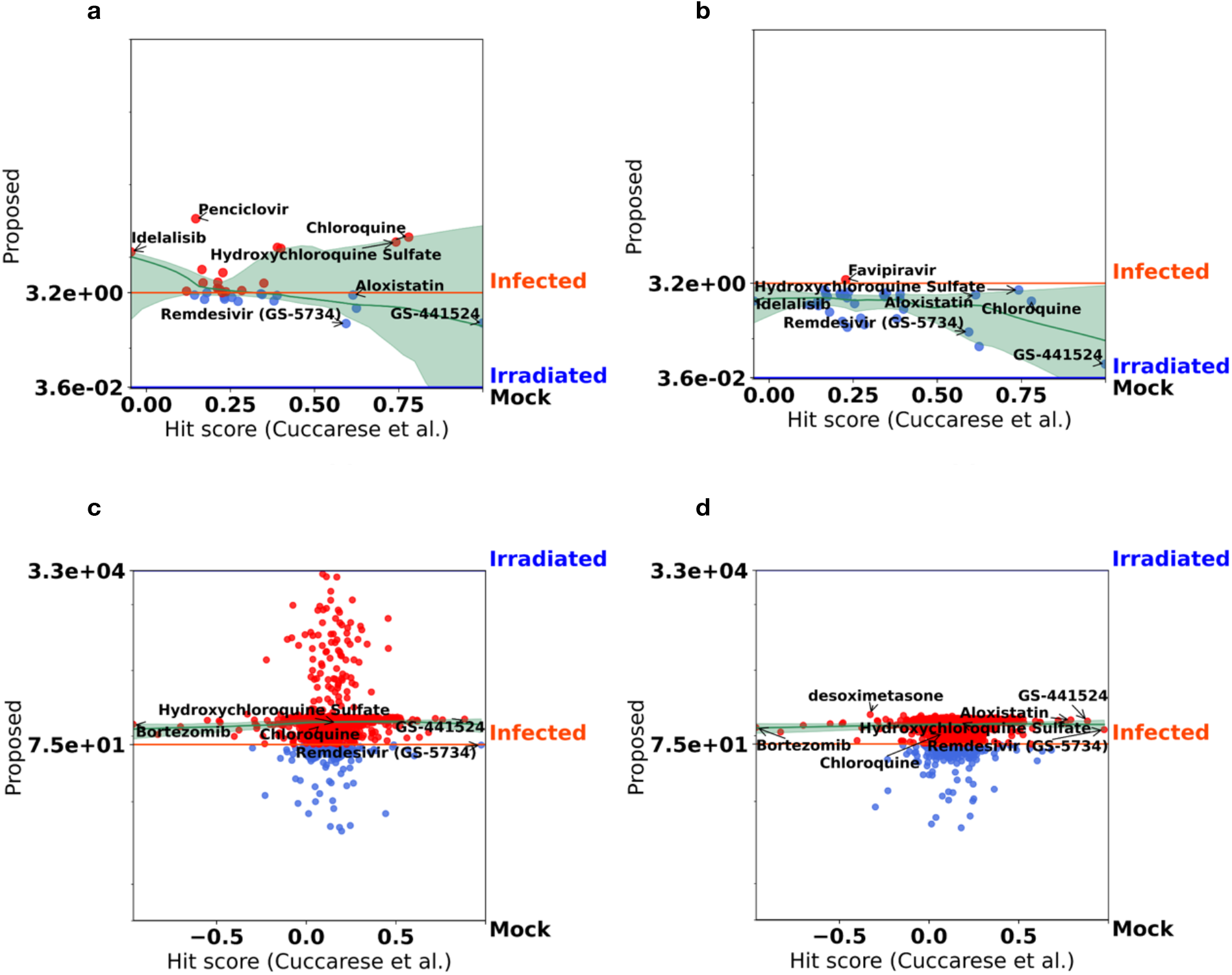
Quantification comparison of cell-based COVID-19 drug responses between the baseline (Cuccarese *et al.* ^5^) and *d*_LEA_ (Ensemble). **a** (VERO) and **c** (HRCE): The quantitative comparison between the hit score^5^ and *d*_LEA_ (Ensemble) with the latent representations of all drug concentrations. **b** (VERO) and **d** (HRCE): The quantitative comparison between the hit score^5^ and *d*_LEA_ with the latent representations of optimal drug concentration.

**Figure 7.**
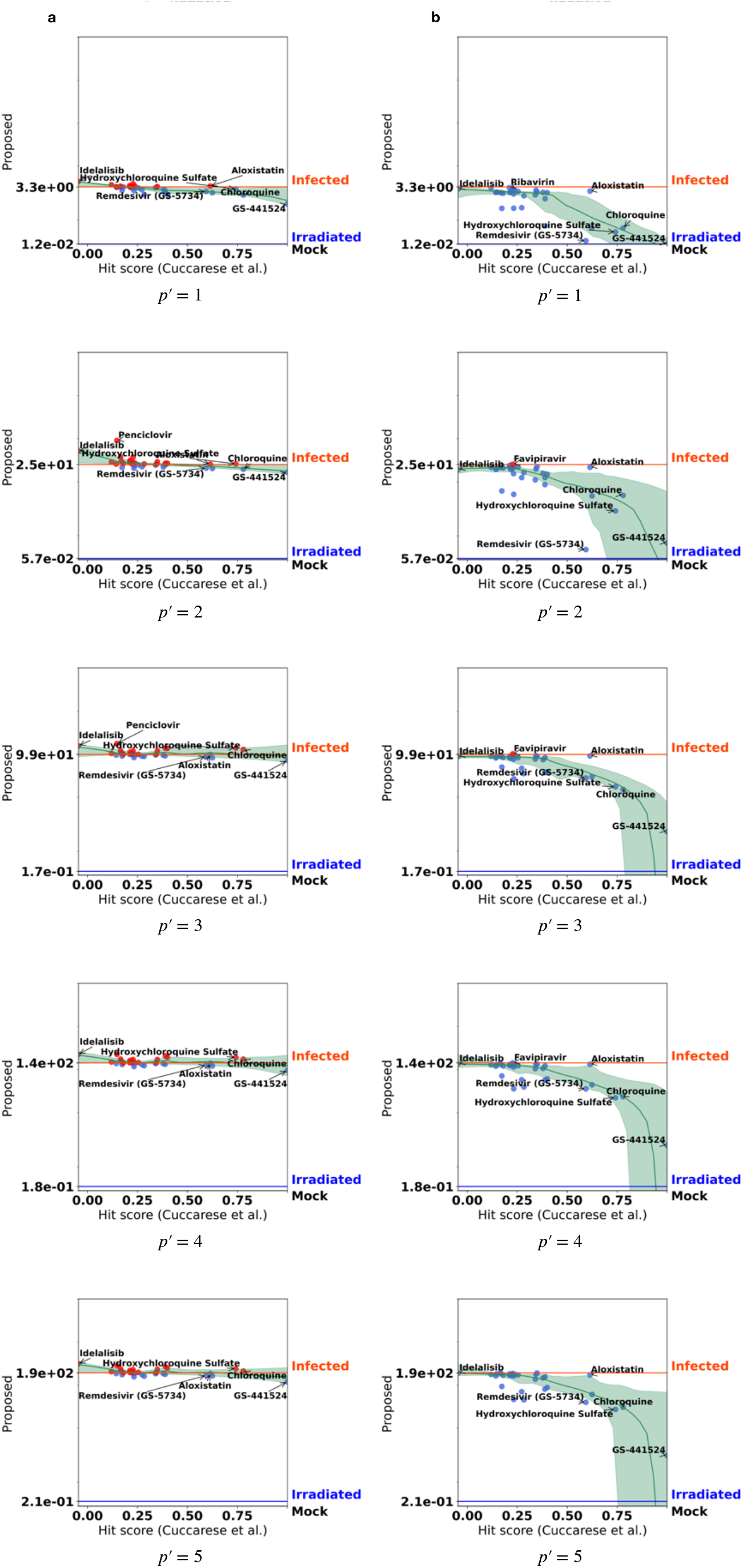
Quantification result *d*_LEA_ of VERO *w.r.t.* different amount of the largest eigenvalues. **a**: *d*_LEA_ computed with the latent representations of all drug concentrations. **b**: *d*_LEA_ computed with the latent representations of optimal drug concentration.

**Figure 8.**
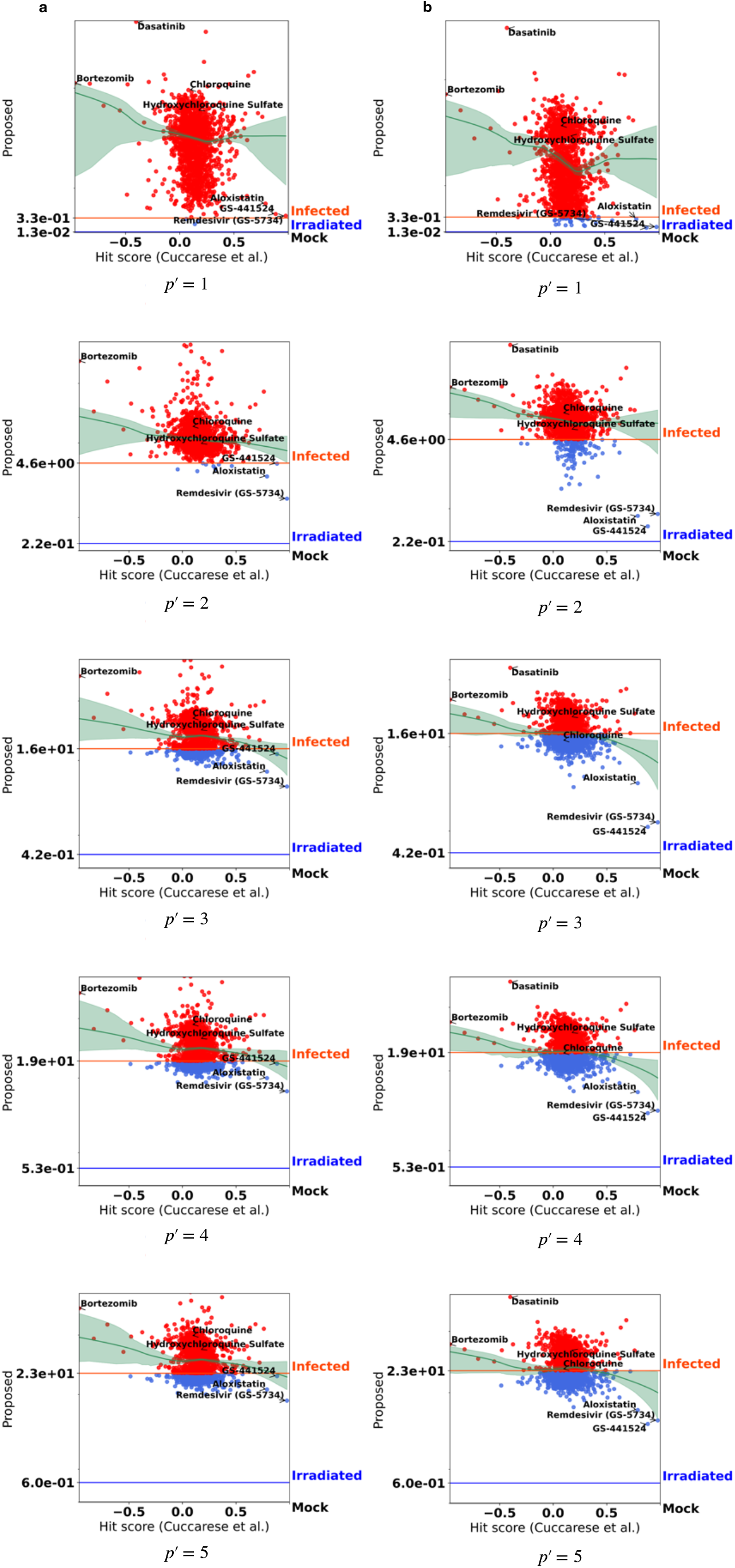
Quantification result *d*_LEA_ of HRCE *w.r.t.* different amount of the largest eigenvalues. **a**: *d*_LEA_ computed with the latent representations of all drug concentrations. **b**: *d*_LEA_ computed with the latent representations of optimal drug concentration.

**Figure 9.**
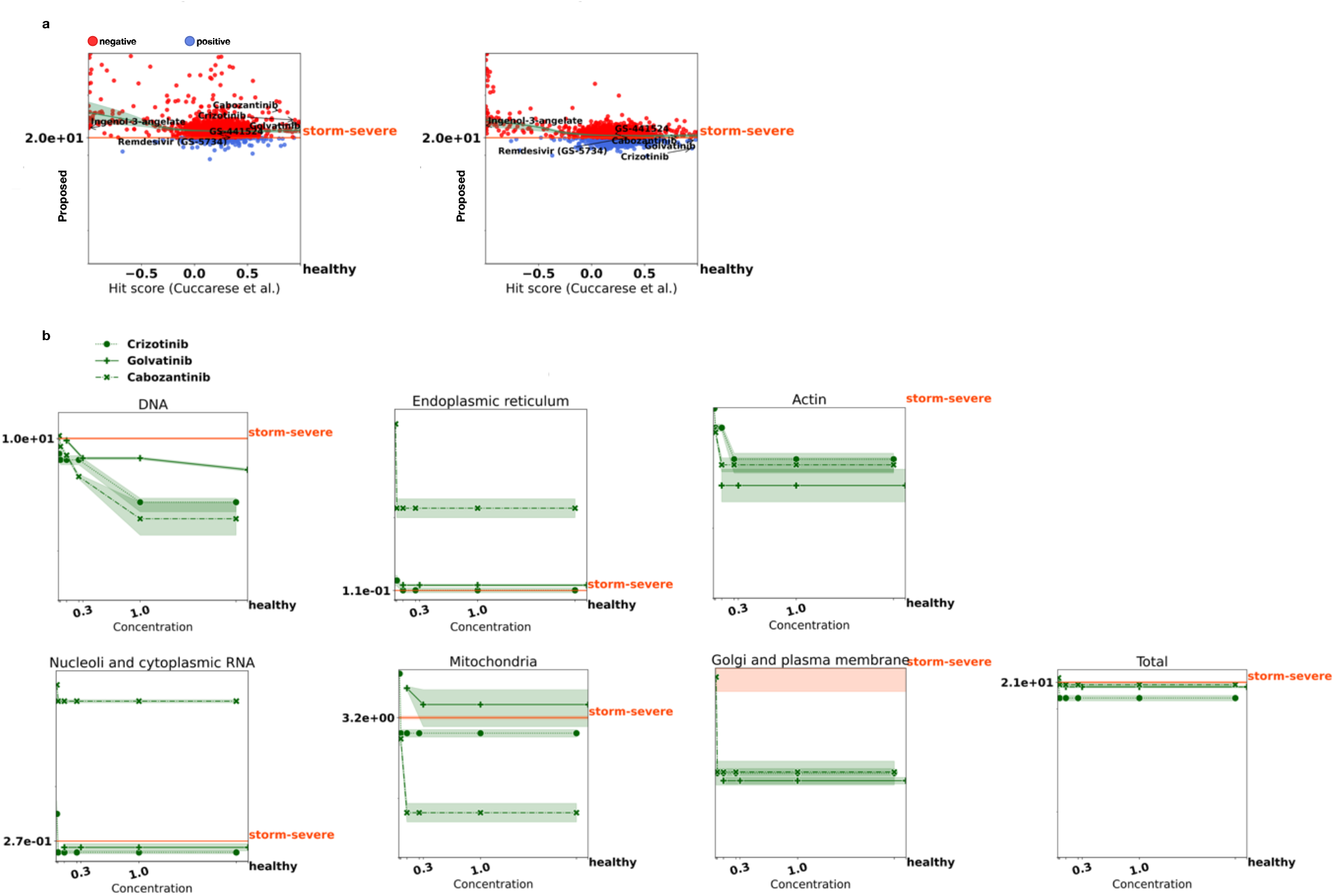
The quantification results of HUVEC experiment. **a**, The quantitative comparison between the hit score^5^ and *d*_LEA_ with latent representations of all drug concentrations (left) and the optimal drug concentration (right). Our drug effects (positive/negative) are thresholded by the *d*_LEA_ of storm-severe cells without drug treatment. **b**, The proposed *d*_LEA_ of different drug concentrations for individual and all fluorescent channels. Here, we report the mean *d*_LEA_ (with standard deviation) averaged on 4 randomly sampled cell collections.

**Figure 10.**
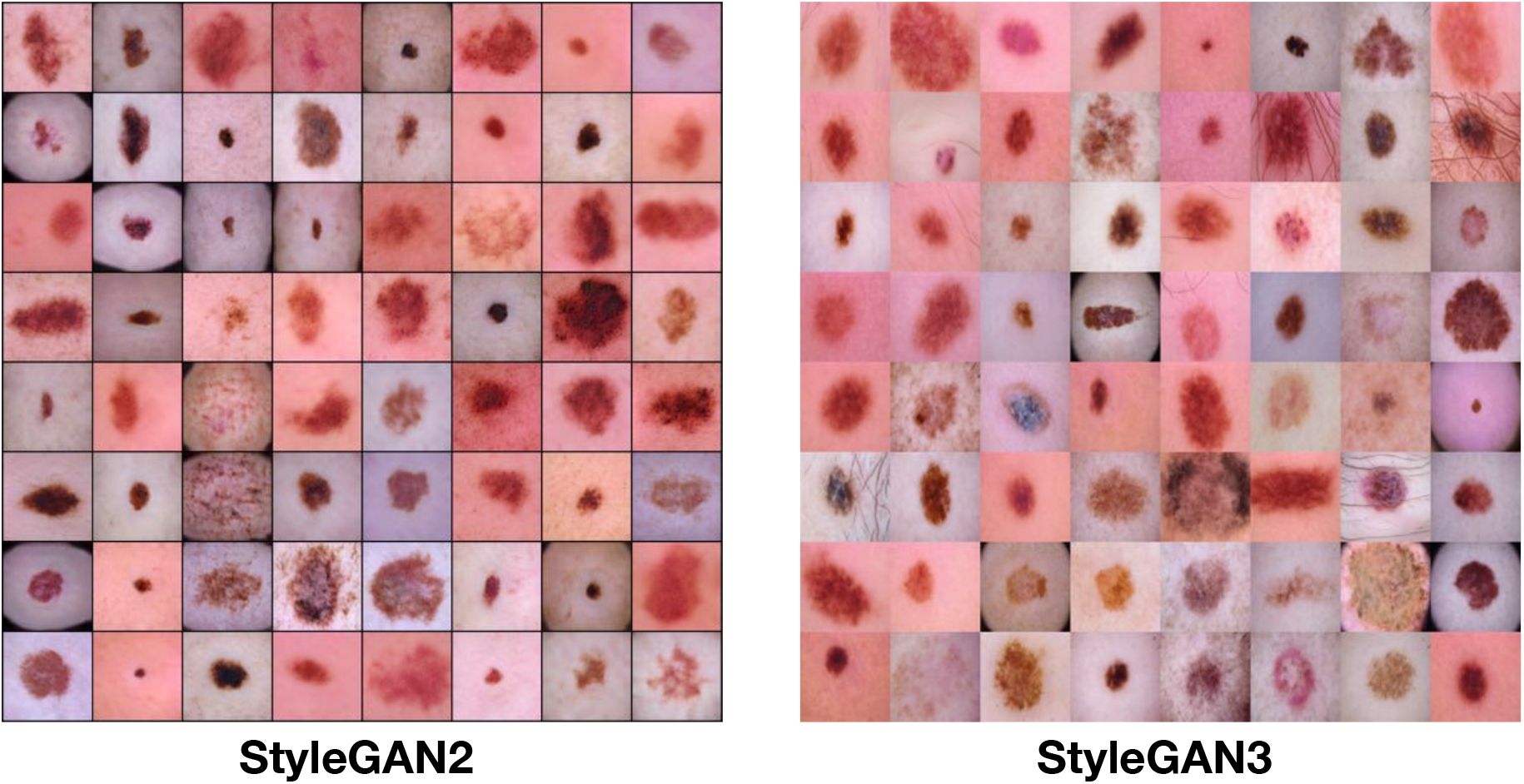
The non-existent skin lesion images synthesized by StyleGAN2 (left) and StyleGAN3 (right).

**Figure 11.**
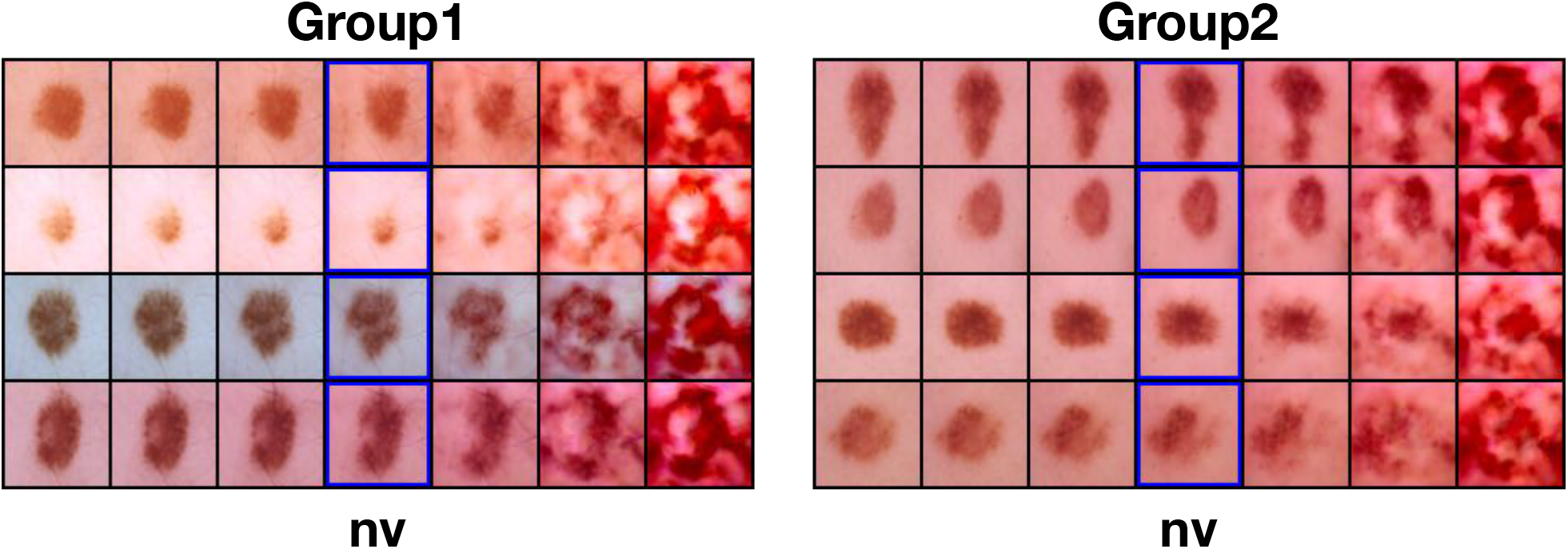
The phenotypic transitions driven by manipulating the largest principal component, which are derived from the latent representations of StyleGAN3_psp.

### Model architecture

Motivated by the impressive achievements of GAN inversion^7^, we instantiate GILEA with the state-of-the-art GAN inversion model (Fig. 1(b)). Firstly, we learn the decoder (generator) under the StyleGAN^28, 29^ framework, which has proved to be successful in hallucinating high-quality natural images. Based on the training protocols suggested in the widely-used repositories^2^ ^3^, we pre-train the StyleGAN2^28^ and StyleGAN3^29^ on HAM10000 to obtain such a decoder (generator) that can synthesize faithful skin lesion images (Please see the Appendix Fig. 10). Next, we launch the encoder training to learn robust latent representations for image reconstruction. Specifically, we apply two practical architectures ‘encoder for editing’^30^ (e4e) and ‘pixel2style2pixel’^31^ (pSp) for comparison, both of which start with a ResNet backbone and then concatenate a feature pyramid network^32^. Similar to the loss design of these studies that enable high-fidelity image reconstruction, we determine our objective to be 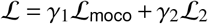, where 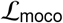 is the contrastive loss that is superior in visual representation learning^33^, and 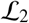 is the *l*_2_ reconstruction loss. Eventually, we report the quantitative reconstruction results in Tab. 2 among compared architectures.

**Table 2.** The numerical comparison of reconstruction results among different model architectures on HAM10000.

| HAM10000 |  | StyleGAN2_pSp (Default) | StyleGAN2_e4e | StyleGAN3_pSp | StyleGAN3_e4e |
| --- | --- | --- | --- | --- | --- |
| nv | PSNR | 30.61 ± 2.20 | 26.40 ± 1.36 | 31.60 ± 2.62 | 30.12 ± 2.44 |
| | SSIM | 0.89 ± 0.06 | 0.85 ± 0.07 | 0.90 ± 0.06 | 0.87 ± 0.06 |
| mel | PSNR | 28.13 ± 2.39 | 24.80 ± 1.69 | 28.60 ± 2.55 | 27.28 ± 2.53 |
| | SSIM | 0.85 ± 0.06 | 0.80 ± 0.08 | 0.86 ± 0.06 | 0.83 ± 0.07 |
| bcc | PSNR | 28.95 ± 2.60 | 25.49 ± 1.58 | 29.48 ± 2.80 | 28.28 ± 2.72 |
| | SSIM | 0.82 ± 0.07 | 0.75 ± 0.08 | 0.83 ± 0.07 | 0.79 ± 0.07 |
| bkl | PSNR | 29.06 ± 2.54 | 25.50 ± 1.48 | 29.55 ± 2.65 | 28.34 ± 2.56 |
| | SSIM | 0.83 ± 0.07 | 0.78 ± 0.08 | 0.85 ± 0.06 | 0.81 ± 0.07 |
| df | PSNR | 30.01 ± 2.29 | 25.96 ± 1.27 | 30.49 ± 2.38 | 29.34 ± 2.37 |
| | SSIM | 0.85 ± 0.06 | 0.79 ± 0.07 | 0.86 ± 0.05 | 0.83 ± 0.06 |
| vasc | PSNR | 30.52 ± 2.77 | 25.78 ± 1.35 | 31.15 ± 3.14 | 29.72 ± 2.92 |
| | SSIM | 0.88 ± 0.07 | 0.84 ± 0.08 | 0.89 ± 0.07 | 0.86 ± 0.08 |
| akiec | PSNR | 27.71 ± 2.09 | 24.86 ± 1.32 | 28.21 ± 2.21 | 27.07 ± 2.12 |
| | SSIM | 0.79 ± 0.06 | 0.71 ± 0.08 | 0.80 ± 0.06 | 0.76 ± 0.07 |
| Total | PSNR | 29.98 ± 2.49 | 26.01 ± 1.54 | 30.81 ± 2.88 | 29.40 ± 2.71 |
| | SSIM | 0.87 ± 0.07 | 0.83 ± 0.08 | 0.88 ± 0.06 | 0.85 ± 0.07 |

**Table 3.** The numerical comparison of reconstruction results among different model architectures on HUVEC.

|  |  | HUVEC (Proposed) | HUVEC (Ensemble) |
| --- | --- | --- | --- |
| DNA | PSNR | $44.70 \pm 3.24$ | $41.03 \pm 3.25$ |
| | SSIM | $0.99 \pm 0.02$ | $0.98 \pm 0.02$ |
| ER | PSNR | $38.11 \pm 3.27$ | $36.90 \pm 3.80$ |
| | SSIM | $0.97 \pm 0.02$ | $0.95 \pm 0.03$ |
| Actin | PSNR | $45.04 \pm 2.68$ | $42.68 \pm 2.72$ |
| | SSIM | $0.98 \pm 0.01$ | $0.97 \pm 0.02$ |
| RNA | PSNR | $42.28 \pm 2.60$ | $41.93 \pm 2.97$ |
| | SSIM | $0.98 \pm 0.01$ | $0.97 \pm 0.02$ |
| Mitochondria | PSNR | $45.11 \pm 2.20$ | $42.93 \pm 2.04$ |
| | SSIM | $0.98 \pm 0.01$ | $0.98 \pm 0.02$ |
| Golgi | PSNR | $42.02 \pm 2.78$ | $39.59 \pm 2.95$ |
| | SSIM | $0.98 \pm 0.01$ | $0.96 \pm 0.03$ |
| Total | PSNR | $41.83 \pm 2.66$ | $40.03 \pm 2.95$ |
| | SSIM | $0.98 \pm 0.01$ | $0.97 \pm 0.02$ |

### pSp VS e4e

In terms of quantitative scores such as PSNR and SSIM, it is clear to see that the pSp encoder in combination with either StyleGAN2 or 3 decoder outperforms e4e by a clear margin, suggesting superior image reconstruction qualities. This can also be verified by the image samples presented in Fig. 5 (a): The images reconstructed from the representations of pSp encoder reserve finer detail of lesion demarcation and skin pigmentation, while the e4e encoder tend to produce more blurry reconstructions. As a result, we take the pSp architecture as the default encoder.

### StyleGAN2 VS StyleGAN3

Following the encoder architecture comparison, we investigate the variants of StyleGAN decoder. While StyleGAN3_pSp achieves better PSNR and SSIM scores than StyleGAN2_pSp, the qualitative difference in image reconstruction appears marginal (Fig. 5 (a)). Furthermore, when examining the *d*_LEA_ behavior with the increasing interpolation weight, we found notable differences between the two architectures. Compared to StyleGAN2, Fig. 5 (b) shows that *d*_LEA_ computed with StyleGAN3_e4e increases unexpectedly from *w* = 0.25 to *w* = 0.5 for the mixture data collection of nv and mel images, which is in conflict with the fact that more inclusion of benign mole images should reduce the distance to the nv reference category. With regards to nv, *d*_LEA_ of StyleGAN3_pSp surprisingly suggests a larger data difference of bkl (bcc) than mel. This is also counter-intuitive as malignant neoplasms (mel) are well known to present distinct appearances in lesion size and pigmentation heterogeneity, allowing these lesions to be clearly differentiated from a benign mole (nv). Besides, we notice that the learned representations of StyleGAN3 tend to be more convoluted and are thus less ideal to support clear biological interpretation (See for example Appendix Fig. 11). Since StyleGAN3 is motivated by the texture-sticking drawback occurring in natural images and imposes equivariant translation and rotation on learned representations^29^, it may explain why StyleGAN3 does not adapt well to biomedical images from substantially distinct modalities. This is also reflected by the drawbacks identified by Alaluf *et al*. ^34^ for natural images.

Based on these results, we set **StyleGAN2_pSp** as the default architecture for conducting drug screening and skin lesion experiments through this article.

### Further comparisons

Next, we investigate the *d*_LEA_ performance in regards to the amount *p*′ of the largest eigenvalues utilized in Eq. 2. As we can see in Fig. 5 (b), *d*_LEA_ shows comparable decreasing scores with the increasing weight of including more nv images for *p*′ = 1,…,4,5. Such results demonstrate the feasibility and robustness of *d*_LEA_ computed with the largest eigenvalues for the RGB imaging. For both the HAM10000 and RxRx19 datasets, we narrow down the amount *p*′ of the largest eigenvalues to 5. In addition, we evaluate *d*_LEA_ using well-established statistical tests and widely used scalar-valued scores. For the former, we report the (average) p-values computed with two collections of eigenvalues. We further compute *d*_KID_ with multiple subsets of randomly sampled latent representations and *d*_Eig_ with two SCMs (Def. 1). Regarding statistical tests, we have witnessed either contradictory behaviors obtained by F_test^35^ or uninformative results from the Levene_test^36^ and Wilcoxon_test^37^, which confirms the challenging adaption of standard statistical tests to high-dimensional use cases. Although meaningful curves illustrating decreasing distances between nv and compared categories can be obtained with *d*_KID_, it shows clear fluctuations with large standard deviations. This is mainly due to the additional randomness that comes from subset selection, which is not present in other measurements. Without the imbalance issue regarding different channels introduced in the skin lesion dataset, *d*_Eig_ shows plausible decreasing trends similar to *d*_LEA_.

### Clinical Interpretations

As shown in Fig. 5 (c), clear patterns in terms of lesion size emerge in two different groups clustered with k-means. This has been persistently presented among different categories. If we manipulate the largest principal component (top rows of Fig. 5 (c)) with increasing 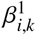 (from left to right), the ‘nv’ images begin to show a poor lesion demarcation, further increase in lesion size and pigmentation heterogeneity, with the lesions displayed towards the right showing clear pathological changes towards a clinically suggestive appearance of malignancy. Considering the largest eigenvalue (×10^5^) 4.41 for nv versus 6.04 for mel, the appearance shift towards malignancy by enlarging the principal component of nv representations can indeed explain the eigenvalue difference 4.41 < 6.04. Comparable observations can be also made when investigating the 5 largest principal components. Apart from similar lesion size patterns arising from the k-means clustering, we show distinct samples clustered in the two groups (Bottom rows of Fig. 5 (c)). Accordingly, the PCA plots regarding the 5 largest eigenvalues verify the distinguished yet consistent heterogeneity quantification among mel, bcc, bkl and nv.

## Author contributions statement

J.W. and V.H.K. conceived the research idea. J.W. implemented the algorithm and carried out the experiments. J.W. and V.H.K. analyzed the results. J.W. and V.H.K. drafted the manuscript. V.H.K. supervised the project.

## Footnotes

1 https://www.fda.gov/drugs/emergency-preparedness-drugs/coronavirus-covid-19-drugs

2 StyleGAN2: https://github.com/rosinality/stylegan2-pytorch.git

3 StyleGAN3: https://github.com/NVlabs/stylegan3.git

## Notes

https://github.com/CTPLab/GILEA

### Competing Interest Statement

The authors have declared no competing interest.

### Summary of Updates

minor revision of the written contents

